# Glioblastoma mutations impair ligand discrimination by EGFR

**DOI:** 10.1101/2021.05.04.442654

**Authors:** Chun Hu, Carlos A. Leche, Anatoly Kiyatkin, Steven E. Stayrook, Kathryn M. Ferguson, Mark A. Lemmon

## Abstract

The epidermal growth factor receptor (EGFR) is frequently mutated in human cancer, and is an important therapeutic target. EGFR inhibitors have been successful in lung cancer, where the intracellular tyrosine kinase domain is mutated, but not in glioblastoma multiforme (GBM) – where mutations (or deletions) occur exclusively in the EGFR extracellular region. Wild-type EGFR is known to elicit distinct signals in response to different growth factor ligands, exhibiting biased agonism. We recently showed that individual ligands stabilize distinct receptor dimer structures, which signal with different kinetics to specify outcome. EGF induces strong symmetric dimers that signal transiently to promote proliferation. Epiregulin (EREG) induces weak asymmetric dimers that generate sustained signaling and differentiation. Intriguingly, several GBM mutation hotspots coincide with residues that define the asymmetric and symmetric dimer structures. Here, we show that common extracellular GBM mutations prevent EGFR from distinguishing between EGF and EREG based on dimer structure and stability – allowing strong dimers to form with both ligands. Crystal structures show that the R84K mutation symmetrizes EREG-driven dimers, whereas the A265V mutation remodels key dimerization sites. Our results suggest that modulating EGFR’s biased agonism plays an important role in GBM, and suggest new approaches for ‘correcting’ aberrant EGFR signaling in cancer.

## INTRODUCTION

In addition to guiding targeted therapy, identification of cancer ‘driver’ missense mutations has yielded important insights into mechanisms of cell signaling (Martínez-Jiménez et al., 2020). For example, the discovery of EGFR mutations in lung cancer catalyzed breakthroughs in understanding how EGFR tyrosine kinase activity is regulated (Zhang et al., 2006) as well as directing use of EGFR-targeted therapeutics (Sharma et al., 2007). Intriguingly, whereas EGFR mutations in lung cancer are largely restricted to the intracellular kinase domain, in GBM they occur almost exclusively in the extracellular region (ECR) (Brennan et al., 2013; Tate et al., 2019). Approximately 24% of GBMs have ECR point mutations in EGFR (Brennan et al., 2013), primarily at ‘hotspots’ marked in Figure 1a (Tate et al., 2019). These mutations occur alongside other *EGFR* aberrations (An et al., 2018a; Brennan et al., 2013), including gene amplification and the common EGFR variant III (vIII) deletion (Sugawa et al., 1990) that removes ECR residues 7 to 273. It remains unclear precisely how *EGFR* aberrations influence GBM, and EGFR inhibitors have not been successful clinically (An et al., 2018a; Eskilsson et al., 2018; Heimberger et al., 2005).

**FIGURE 1.**
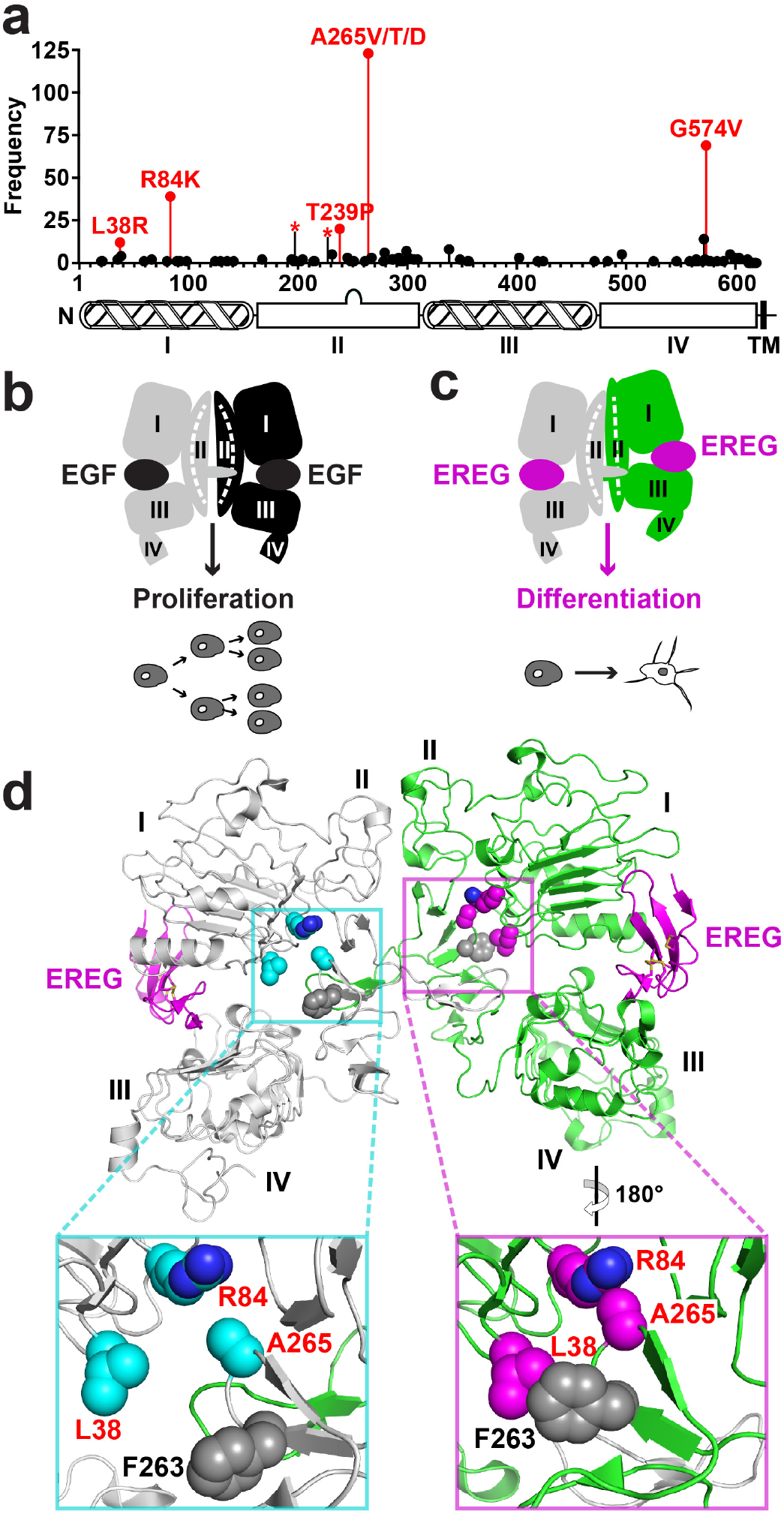
Extracellular EGFR mutations in glioblastoma, and location in structure. **a.** Distribution of mutations in the extracellular region of EGFR found in GBM in the COSMIC database (Tate et al., 2019), with the most common non-cysteine mutations colored red and frequency (number of occurrences in COSMIC (Tate et al., 2019)) depicted by bar height. R198C and R228C are represented as red asterisks. Residue numbering corresponds to the mature form of EGFR, subtracting 24 for the signal sequence (R84 is R108 in pro-EGFR and A265 is A289). Below the graph is a schematic of the EGFR extracellular region domain structure. **b.** Cartoon of a symmetric dimer of the EGFR extracellular region (sEGFR) induced by EGF (black), with associated proliferative response in MCF7 cells (Freed et al., 2017). Individual domains are numbered. **c.** Cartoon of the asymmetric wild-type sEGFR dimer induced by EREG (magenta), with associated differentiative response in MCF7 cells (Freed et al., 2017). **d.** Structure of the EREG-induced asymmetric wild-type sEGFR dimer (PDBID 5WB7) (Freed et al., 2017). The left-hand molecule (grey) resembles sEGFR in symmetric ligand-bound sEGFR dimers, with a ligand-induced bend in domain II (see curve in domain II cartoon in left-hand side of **c**). The right-hand molecule (light green) does not have the ligand-induced bend in domain II (see straight line in right-hand molecule in **c**). Key residues mutated in GBM are colored cyan (left) or magenta (right), and are labelled in the inserts, as is F263. Note that the right-hand insert is rotated 180° about a vertical axis to allow comparison with L38/R84/A265 positions in the lefthand molecule.

Individual ligands are known to induce distinct cell fates through EGFR (Wilson et al., 2009), in some cases by stabilizing receptor dimers with different structures and stabilities (Freed et al., 2017) in a form of biased agonism (Macdonald-Obermann and Pike, 2014). EGF stabilizes symmetric, strong dimers of the ECR (Figure 1b), driving transient signaling and proliferation. EREG induces much weaker asymmetric dimers (Figure 1c), which promote sustained signaling and differentiation in some cell lines (Freed et al., 2017). Importantly, three of the most frequently mutated EGFR residues in GBM (L38, R84 and A265: numbered for mature EGFR) (Binder et al., 2018; Brennan et al., 2013; Tate et al., 2019) cluster in the 3-dimensional structure (Ferguson et al., 2003; Ng et al., 2018) (Figure 1d) and play a key role in defining the asymmetry of EREG-induced sEGFR dimers. In the left-hand molecule in Figure 1d (and in EGF-induced sEGFR dimers) these three residues are well separated from one another. By contrast, in the right-hand EGFR molecule of this asymmetric dimer, R84 and A265 contact one another, and L38 makes van der Waal’s contacts with F263 (Figure 1d, insert). Identical interactions are also seen between domains I and II of unliganded monomeric EGFR (Ferguson et al., 2003). Since EREG is highly expressed in GBM, and its expression is associated with reduced survival (Auf et al., 2013; Kohsaka et al., 2014; Shergalis et al., 2018; von Achenbach et al., 2018), we hypothesized that mutating these residues might symmetrize and strengthen EREG-induced EGFR dimers. This could ‘switch’ the EGFR response to EREG binding from one that promotes differentiation to one that is ‘EGF-like’, with EREG inducing strong receptor dimers that promote proliferation. The resulting loss of EREG-induced differentiation (or enhanced proliferation) might contribute to tumor formation (Friedmann-Morvinski et al., 2012; Zheng et al., 2010).

We show here that common extracellular GBM mutations prevent EGFR from distinguishing between EGF and EREG based on dimer structure and stability – allowing strong dimers to form with both ligands. Crystal structures show that the R84K mutation symmetrizes EREG-driven dimers, whereas the A265V mutation remodels key dimerization sites. Our results suggest that modulating EGFR’s biased agonism plays an important role in GBM, and may suggest new approaches for ‘correcting’ aberrant EGFR signaling in cancer.

## RESULTS AND DISCUSSION

### GBM mutations selectively enhance EREG-induced cell proliferation

Several studies have shown that EGFR variants harboring GBM mutations can promote IL-3-independent growth of Ba/F3 cells (Kohsaka et al., 2017; Lee et al., 2006; Ng et al., 2018) – a sign of transforming ability (see also Figure 2a). When assayed in the presence of serum, EGFR variants harboring GBM mutations have also been reported to enhance anchorage-independent growth, tumorigenicity, or invasion of NIH 3T3 or U87 cells (Binder et al., 2018; Lee et al., 2006). Although this has led them to be classified as driver mutations (Ng et al., 2018), biochemical studies show that these variants only become substantially activated when ligand is added (Binder et al., 2018; Lee et al., 2006) (see also Figure 2b). Moreover, we showed in previous work that the common GBM-associated ECR mutations do not promote ligand-independent dimerization of the EGFR ECR (Bessman et al., 2014a), although they do enhance ligand-binding affinity. For at least some GBM mutations, we found in Ba/F3 cell studies that the major effect may be sensitization of EGFR to certain ligands – particularly with the R84K mutation, which had a disproportionately large effect on EREG-induced proliferation (Figure 2c). Thus, GBM mutations in EGFR do not seem to cause strong constitutive activation *per se*, but may alter relative signaling responses to the different ligands in serum.

**FIGURE 2.**
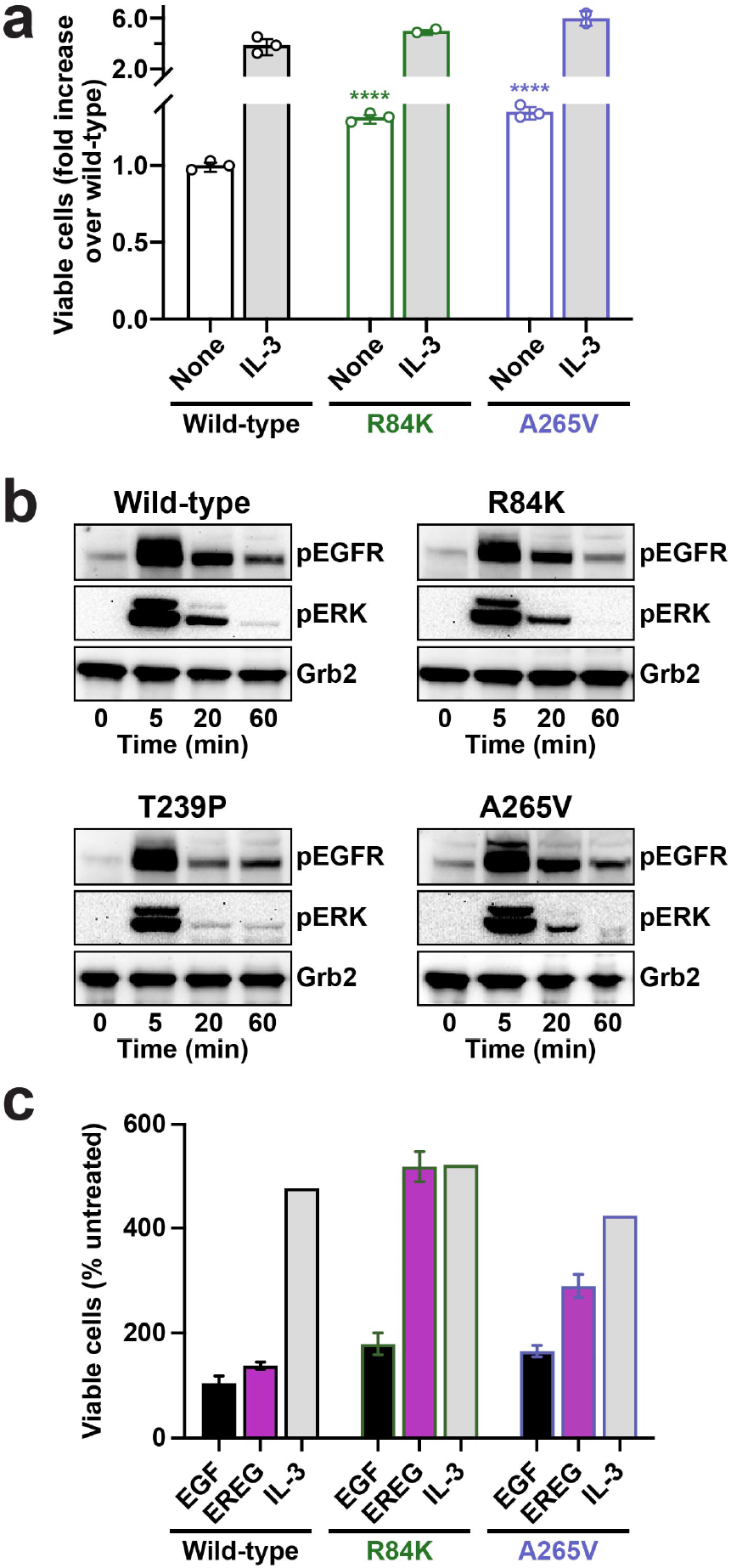
Ligand-dependence of GBM-mutated EGFR, and differential effects of GBM mutations on EREG and EGF signalling. **a.** IL-3-dependent Ba/F3 cells were stably transfected with human wild-type EGFR or with variants harboring an R84K or A265V mutation. Cells were either left untreated or were treated with IL-3 (2 ng/ml) for 72 h, after which a CyQuant Direct proliferation assay was used to detect the number of viable cells in each condition, and resulting fluorescence signals were normalized to that seen with untreated wild-type and shown as mean ± SD (n = 3 experiments). IL-3 treatment promotes robust proliferation in all cases (as positive control). As previously reported (Kohsaka et al., 2017; Lee et al., 2006; Ng et al., 2018), the mutated EGFRs promoted statistically significant increases in viable cell numbers compared with wild-type EGFR in the absence of ligand, but the effects were small at these expression levels – with the number of viable cells increased (compared with wild-type) by 1.31-fold (p = 0.0002) for R84K and 1.35-fold for A265V (p = 0.0003). p values are for unpaired two-tailed Student’s t-tests. **b.** Full-length human EGFR – wild-type or harboring an R84K, T239P or A265V mutation – was stably expressed in the engineered haploid eHAP cell line (Essletzbichler et al., 2014), which has negligible levels of endogenous EGFR (undetectable by Western blotting). Stably transfected cells were serum-starved overnight and either left unstimulated or stimulated with EGF (100 ng/ml) for the indicated times. Levels of phosphorylated EGFR (pY845: CST #2231) and ERK1/2 (pT202/pY204: CST #9106), were then detected by immunoblotting of total cell lysates, with blotting for Grb2 used as loading control (Aksamitiene et al., 2015; Freed et al., 2017). Both EGFR phosphorylation and ERK phosphorylation are ligand-dependent in all cases, with no evidence for constitutive activation of the mutated receptors. **c.** Ba/F3 cells stably transfected with wild-type or mutated EGFR were sorted by flow cytometry to yield cell populations expressing similar levels of EGFR at their surface. Cells were either left untreated or were treated with similarly saturating levels of EGF (2 nM) or EREG (100 nM) for 72 hours. The CyQuant Direct proliferation assay was used to detect the number of viable cells in each condition. Fluorescence signals were normalized by signals from untreated cells and are shown as mean ± SD (n = 3 experiments). With equal receptor expression levels, A265V-mutated EGFR showed no significant differences compared with wild-type EGFR. However, R84K and A265V showed particularly large increases in EREG-induced proliferation compared to wild-type.

### GBM mutations stabilize EREG-induced EGFR dimers

To test the hypothesis that GBM mutations strengthen EREG-induced EGFR dimers, we measured dimerization of the EREG-bound purified recombinant ECR (residues 1-501) from EGFR (sEGFR). The L38R, R84K, A265V, and A265T mutations all enhanced EREG-induced sEGFR dimerization by at least 100-fold. The enhanced dimerization was readily seen in small-angle X-ray scattering (SAXS) studies (Figure 3), which give quantitative shape-independent molecular weight information for proteins in solution (based on *y* intercept, or *I(0)* in Guinier plots shown in Figure 3a-e). Wild-type sEGFR is completely monomeric at 70 μM, but adding EGF doubles the *I*(0) value (based on the *y* intercept in Figure 3a) – representing complete sEGFR dimerization (Lemmon et al., 1997). This is consistent with the dissociation constant (*K*_d_) measured for dimers of EGF-bound sEGFR (1-3 μM) (Dawson et al., 2005; Lemmon et al., 1997). By contrast, EREG failed to induce wild-type sEGFR dimerization in this experiment, giving the same *y* intercept as unliganded sEGFR (Figure 3a) (Freed et al., 2017). Remarkably, all of the GBM mutations allowed EREG to induce almost complete sEGFR dimerization in identical SAXS experiments, but did not greatly impact ligand-independent sEGFR dimerization (Figures 3b-e). The enhanced EREG-induced dimerization of GBM-mutated sEGFR was also evident in chemical crosslinking studies (Figures 3a-e, lower panels and Figure S1). To go from undetectable EREG-induced dimerization to complete dimerization at 70 μM sEGFR (Figure 3f) implies that the *K*_d_ value for dimerization of the EREG:sEGFR complex has been reduced from >700 μM (wild-type) to <7 μM by these mutations – *i.e*. is enhanced by at least 100-fold.

**FIGURE 3.**
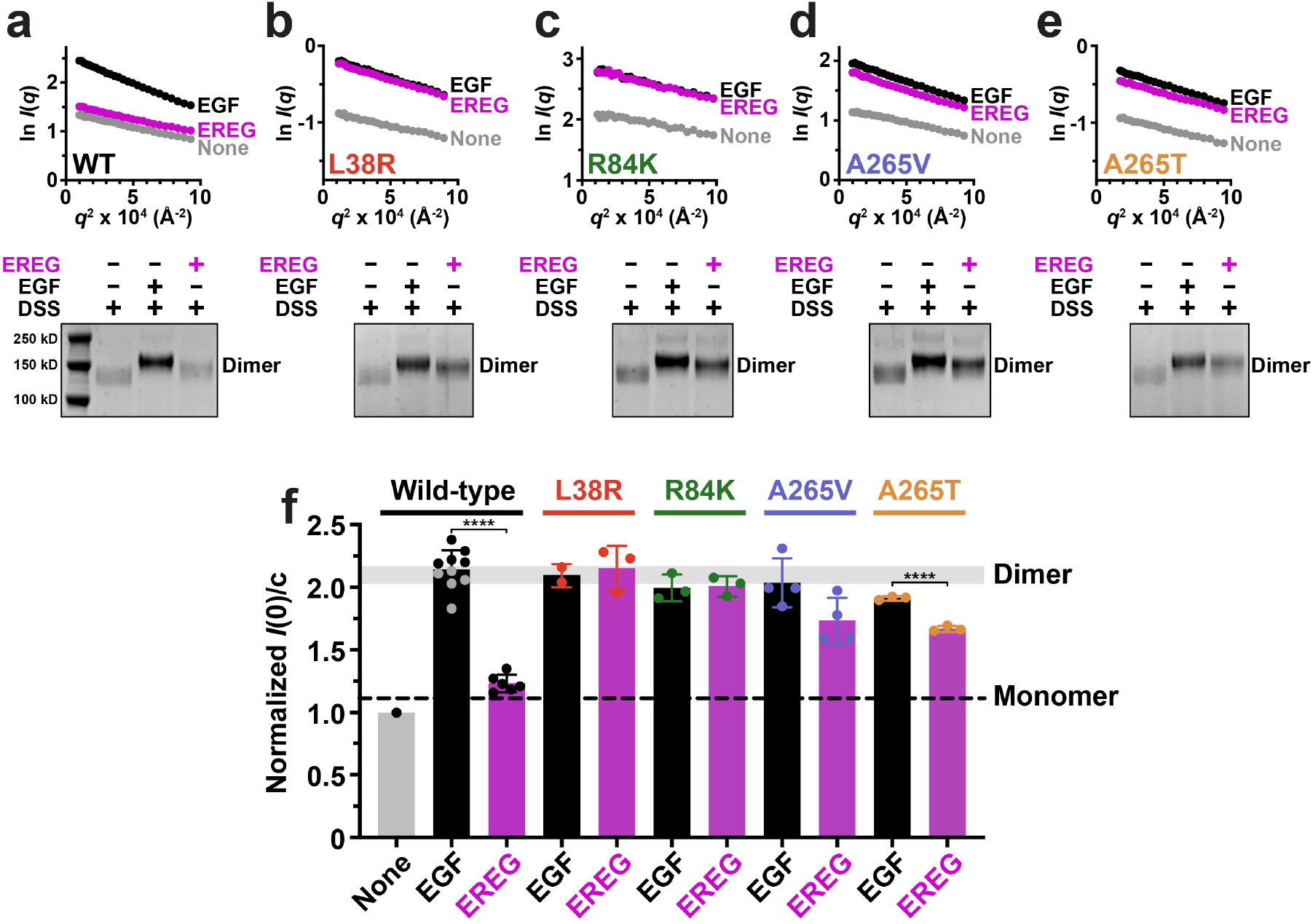
Glioblastoma mutations selectively enhance EREG-induced EGFR dimerization. **a.-e.** Small-angle X-ray scattering and chemical cross-linking studies of ligand-induced sEGFR dimerization for wild-type sEGFR and sEGFR harboring GBM mutations. Upper panels show Guinier plots of SAXS data is shown. The natural logarithm of the scattering intensity at a particular angle, *I*(*q*), normalized for mass concentration, is plotted against *q*^2^ (see Experimental Procedures) and extrapolated to the *y* axis intercept to estimate *I*(0), which is proportional to the weight-averaged molecular mass of molecules in a SAXS sample: ln *I*(*q*) = ln *I*(0) – (*R*_g_^2^/3)*q*^2^, where *R*_g_ is the radius of gyration. For wild-type sEGFR, *I*(0) doubles upon EGF binding (**a**), representing dimerization, but is unaffected by EREG – which fails to induce sEGFR dimers at this concentration (70 μM)(Freed et al., 2017). By contrast, for sEGFR harboring an L38R (**b**), R84K (**c**), A265V (**d**), or A265T (**e**) mutation, EREG increases *I*(0) to a similar extent as EGF, reflecting strong EREG-induced sEGFR dimerization. Lower panels show chemical cross-linking experiments (see Experimental Procedures) illustrating strong EGF-induced dimerization of all sEGFR variants at 5 μM, and the ability of each GBM mutation to enhance EREG-induced sEGFR dimerization (minimal with wild-type). Full gels are shown in Figure S1, with quantitation of cross-linking data. **f.** Summary of SAXS *I*(0) measurements across multiple repeats. For wild-type sEGFR, only EGF (black) doubles *I*(0), representing selective EGF-induced dimerization. By contrast, both EREG (magenta) and EGF (black) induce dimerization of sEGFR harboring L38R (red), R84K (green), A265V (blue) or A265T (gold) mutations – with EREG-induced dimerization of A265 variants appearing slightly less robust. Data represent mean *I*(0)/c ± SD for at least three biological repeats. The degree of sEGFR dimerization for EGF and EREG is significantly different only for wild-type (p < 0.0001) and A265T (p < 0.0001). p values are for unpaired two-tailed Student’s t-tests. See also Figure S1.

Importantly, the effect seen here is ligand-specific, with GBM mutations selectively strengthening dimers induced by EREG. By contrast, the strength of sEGFR dimers induced by transforming growth factor-α (TGFα) was not significantly affected by L38R or R84K mutations in sedimentation equilibrium analytical ultracentrifugation (SE-AUC) experiments (Figure S2a), and the A265V or A265T mutations actually slightly weakened TGFα-induced dimers. This ligand-dependent effect is also seen only for extracellular mutations found in GBM. A recently-described rare extracellular EGFR lung cancer mutation, M253E (Yu et al., 2017)) instead enhanced both ligand-independent and ligand-induced sEGFR dimerization (Figure S2b). Thus, the GBM mutations highlighted in Figure 1a appear to selectively enhance the ability of only certain ligands (e.g. EREG) to induce EGFR dimerization, arguing that they alter the ability of EGFR to distinguish between its activating ligands.

### R84K mutation ‘symmetrizes’ EREG-induced sEGFR dimers

We next used X-ray crystallography to ask how GBM mutations specifically enhance EREG-induced sEGFR dimerization. We first determined the structure of an EREG-induced dimer of R84K-mutated sEGFR to 2.9 Å resolution (Figures 4 and 5a; Table 1). Remarkably, this structure confirmed the hypothesis that mutating R84 to lysine symmetrizes EREG-induced sEGFR dimers. The resulting dimer closely resembles the strong symmetric wild-type sEGFR dimer induced by TGFα (Garrett et al., 2002) or EGF (Lu et al., 2010; Ogiso et al., 2002) (Figure 4).

**FIGURE 4.**
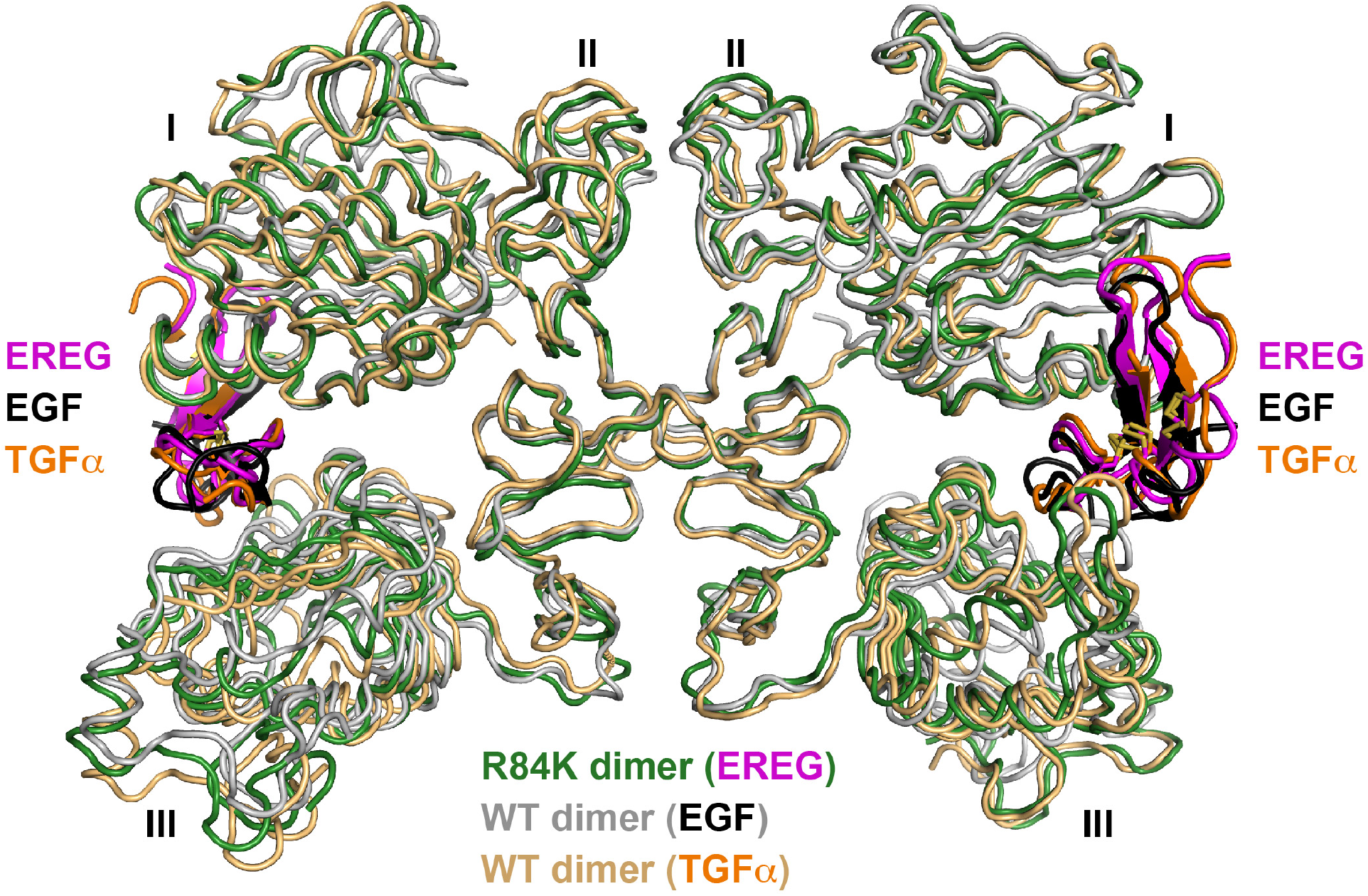
Symmetry of the EREG-induced R84K sEGFR dimer. **a.** Overlay of the EREG-induced R84K-mutated sEGFR dimer (dark green ribbons) with the symmetric dimers of wild-type sEGFR induced by TGFα (PDBID 1MOX) (Garrett et al., 2002) in gold ribbons or EGF (PDBID 3NJP) (Lu et al., 2010; Ogiso et al., 2002), as grey ribbons. EREG, TGFα and EGF are colored magenta, orange, and black respectively.

**FIGURE 5.**
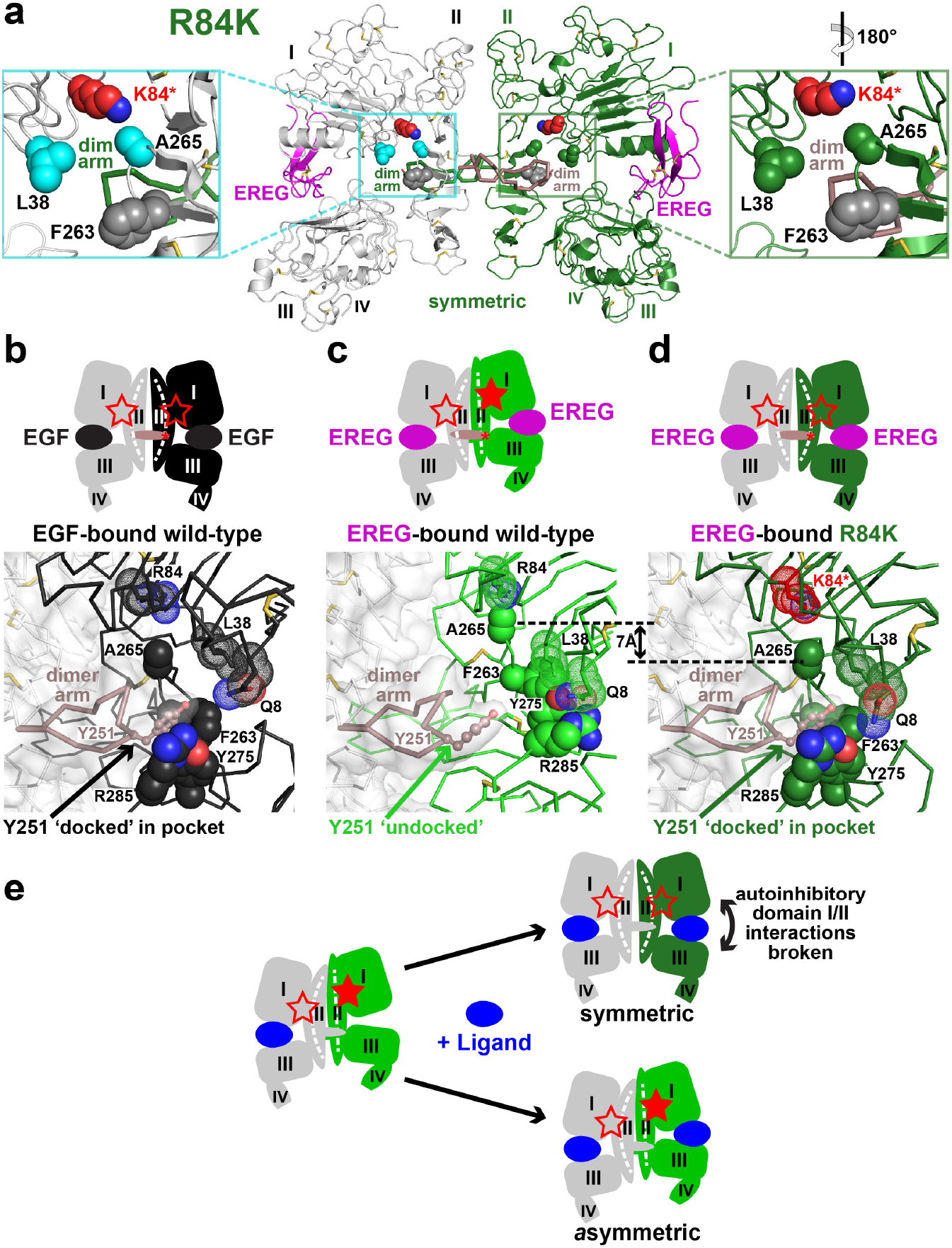
The R84K GBM mutation symmetrizes EREG-induced EGFR dimers. **a.** Cartoon structure of EREG-induced dimer of R84K-mutated sEGFR, with left-hand molecule colored grey and the right-hand molecule colored dark green. The structure is symmetric, and autoinhibitory interactions involving A265 and L38 (cyan in left molecule, green in right) are effectively disrupted, as shown in zoomed regions (note 180° rotation of right-hand zoom to allow structural comparison with left-hand zoom). EREG is shown in magenta, and the side-chain of the K84 substitution in red. Dimer arms are marked, and shown as ribbons. **b.** Upper panel: Cartoon of EGF-induced symmetric (wild-type) sEGFR dimer, emphasizing that the L38/F263 and R84/A265 autoinhibitory interactions are broken in both molecules (open red stars). The dimer arm is colored salmon, with Y251 at its tip represented by a red asterisk. Lower panel: Close-up of the Y251 side-chain in the dimer arm of the left-hand molecule of an EGF-induced wild-type sEGFR dimer docked into its binding site formed by F263, Y275, and R285 from the right-hand molecule. Key side-chains from domain I are shown as black dotted spheres (colored by atom), and those from domain II are shown as solid black spheres. The dimer arm is salmon, and the Y251 side-chain is shown. **c.** Upper panel: Cartoon of EREG-induced asymmetric (wild-type) sEGFR dimer, emphasizing that the L38/F263 and R84/A265 autoinhibitory interactions are broken in the left-hand molecule (open red stars) but not the right-hand molecule (filled red star). Lower panel: Close-up, showing failure of the dimer arm Y251 side-chain to dock onto the righthand molecule (light green) in the EREG-induced wild-type sEGFR dimer. As depicted by the filled red star in the right-hand molecule of the upper panel, A265 and R84 retain their autoinhibitory interaction, as do L38 and F263. This distorts the F263/Y275/R285 dimer arm docking site, so that Y251 fails to dock. Key side-chains from domain I are shown as light green dotted spheres (colored by atom), and those from domain II are shown as solid light green spheres. Dimer arm shown as in **c**. **d.** Upper panel: Cartoon of EREG-induced asymmetric (R84K) sEGFR dimer, emphasizing that the L38/F263 and R84/A265 autoinhibitory interactions are broken in both molecules (open red stars) in the symmetric dimer formed by this variant. Lower panel: Close-up view for the EREG-induced R84K dimer showing ‘release’ of A265 from R84 when it is mutated to K (red), allowing domain II to shift by ~7 Å (shown) – which permits the F263/Y275/R285 dimer arm docking site to form and to accommodate Y251 from the opposing molecule. Key side-chains from domain I are shown as dark green dotted spheres (colored by atom), and those from domain II are shown as solid dark green spheres. Dimer arm shown as in **c**. The 7 Å shift of A265 (and residues C-terminal to is) between wild-type and R84K is shown. **e.** Schematic of negative cooperativity in ligand binding to EGF receptors (Alvarado et al., 2010; Bessman et al., 2014b; Liu et al., 2012) for any ligand (blue) as described in text. Open and closed red stars depict loss and maintenance respectively of autoinhibitory interactions involving L38, R84, F263, and A265 (see also Figure S3). See also Figure S2.

**Table 1.**
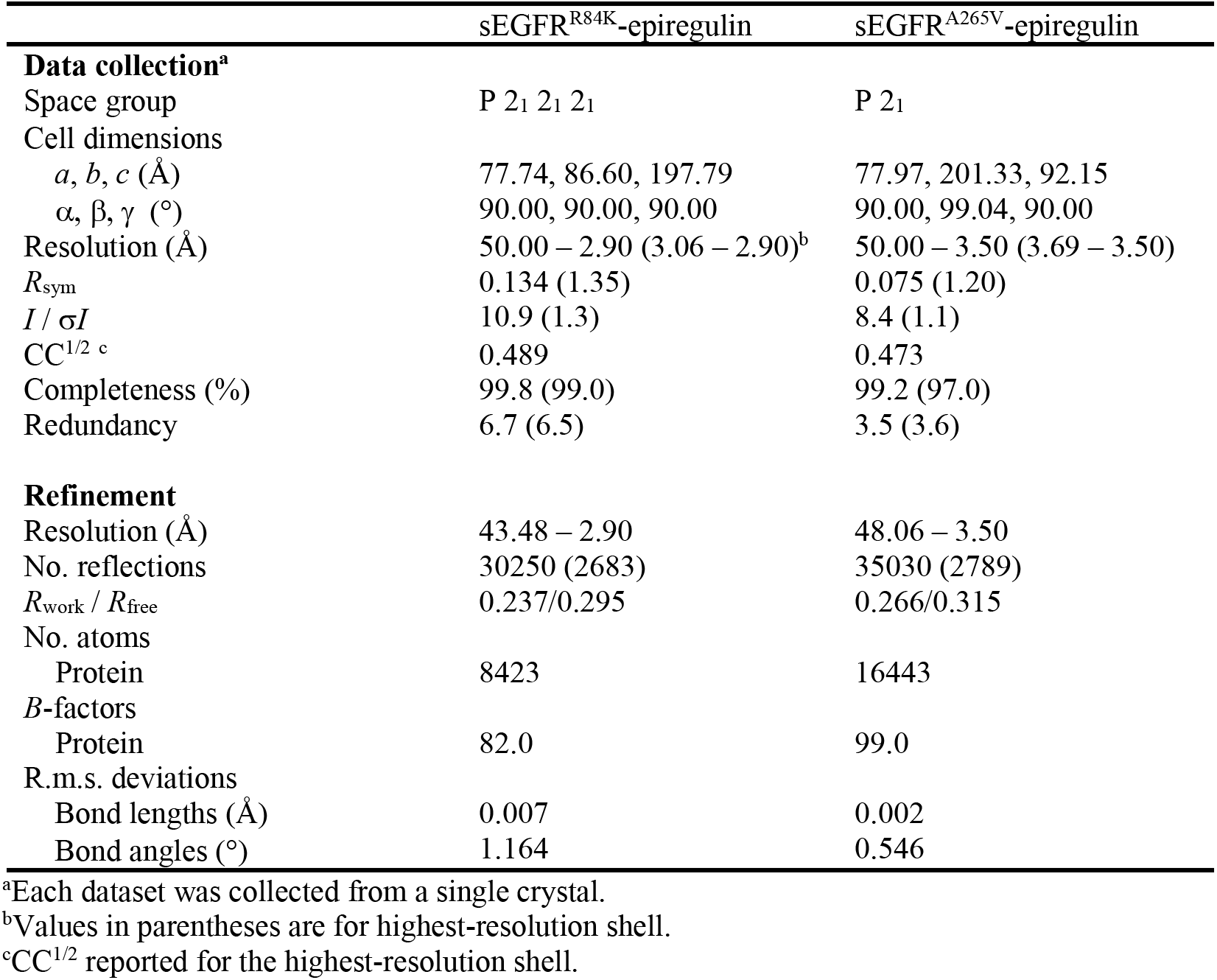
Data collection and refinement statistics

|  | sEGFR<sup>R84K</sup>-epiregulin | sEGFR<sup>A265V</sup>-epiregulin |
| --- | --- | --- |
| **Data collection<sup>a</sup>** |  |  |
| Space group | P 2<sub>1</sub> 2<sub>1</sub> 2<sub>1</sub> | P 2<sub>1</sub> |
| Cell dimensions |  |  |
| *a*, *b*, *c* (Å) | 77.74, 86.60, 197.79 | 77.97, 201.33, 92.15 |
| $\alpha$, $\beta$, $\gamma$ (°) | 90.00, 90.00, 90.00 | 90.00, 99.04, 90.00 |
| Resolution (Å) | 50.00 – 2.90 (3.06 – 2.90)<sup>b</sup> | 50.00 – 3.50 (3.69 – 3.50) |
| *R*<sub>sym</sub> | 0.134 (1.35) | 0.075 (1.20) |
| *I* / σ*I* | 10.9 (1.3) | 8.4 (1.1) |
| CC<sup>1/2</sup> <sup>c</sup> | 0.489 | 0.473 |
| Completeness (%) | 99.8 (99.0) | 99.2 (97.0) |
| Redundancy | 6.7 (6.5) | 3.5 (3.6) |
| **Refinement** |  |  |
| Resolution (Å) | 43.48 – 2.90 | 48.06 – 3.50 |
| No. reflections | 30250 (2683) | 35030 (2789) |
| *R*<sub>work</sub> / *R*<sub>free</sub> | 0.237/0.295 | 0.266/0.315 |
| No. atoms |  |  |
| Protein | 8423 | 16443 |
| *B*-factors |  |  |
| Protein | 82.0 | 99.0 |
| R.m.s. deviations |  |  |
| Bond lengths (Å) | 0.007 | 0.002 |
| Bond angles (°) | 1.164 | 0.546 |

<sup>a</sup>Each dataset was collected from a single crystal.<sup>b</sup>Values in parentheses are for highest-resolution shell.<sup>c</sup>CC<sup>1/2</sup> reported for the highest-resolution shell.

In monomeric sEGFR without bound ligand (or with epigen bound), the GBM mutation hot-spot residues L38, R84, and A265 are engaged in direct (R84/A265 and L38/F263) side-chain contacts (Ferguson et al., 2003), which restrain the relative positions of domains I and II (Figure S3a,b). These contacts are fully broken in symmetric wild-type sEGFR dimers induced by TGFα (Garrett et al., 2002) or EGF (Lu et al., 2010; Ogiso et al., 2002) (Figure S3c), as depicted by open red stars in Figure 5b (upper cartoon). In the EREG-induced asymmetric dimer, by contrast, one set remains unbroken (Figure 1d, and Figure S3b, lower), as noted with a filled star in the upper cartoon of Figure 5c. The R84K mutation allows EREG to break these contacts in both protomers of the dimer (Figure 5a,d). This is the root cause of both the symmetrization and the enhanced EREG-induced dimer strength that result from this mutation, as described below.

The dimer arm plays a key role in stabilizing EGF-induced sEGFR dimers (Figure 5b), with the side-chain at its tip (from Y251) docking into a shallow pocket formed by F263, Y275 and R285 in the opposing molecule (right-hand in Figure 5b). Indeed, mutating Y251 and R285 completely abolishes EGFR dimerization without preventing ligand binding (Dawson et al., 2005; Ogiso et al., 2002). In order for the F263/Y275/R285 pocket to form in the location required for dimer arm docking, the L38/F263 and R84/A265 side-chain interactions must be broken in the right-hand molecule of Figure 5b – as well as the left-hand molecule. In the asymmetric EREG-induced wild-type sEGFR dimer (Freed et al., 2017) (Figure 5c), by contrast, these autoinhibitory interactions are not broken in the right-hand molecule, and domain II is kept ‘fixed’ against domain I. This prevents the bend (around residue 238) seen in domain II when sEGFR dimerization is induced by EGF or TGFα (Figure S3d) (Ferguson, 2008). F263, Y275 and R285 are thus held away from the dimer interface (through interactions with Q8 and L38 in domain I), and fail to form the F263/Y275/R285 pocket that optimally accommodates Y251. The side-chain of Y251 (and consequently the dimer arm) thus remains undocked (Figure 5c) – explaining the weak dimerization.

Mutating R84 to lysine (red in Figures 5a and d) ‘releases’ A265 so that domain I/II interactions are broken and domain II can bend – as seen by the >7Å movement of A265 down in the plane of the page when comparing lower panels of Figures 5c and d. The L38/F263 interaction is also broken (see open star in upper cartoon in Figure 5c), freeing the side-chains of F263, Y275 and R285 to move toward the dimer interface and to reform the (EGF dimer-like) docking site for Y251 on the opposing dimer arm (Figure 5d). These changes allow stabilization of the EREG-induced sEGFR dimer as seen in our SAXS studies.

### R84K mutation equalizes EREG binding sites

In addition to strengthening sEGFR dimerization by symmetrizing the dimer interface, the R84K mutation permits the two ligand binding sites in the dimer to be highly similar (Figure 6a) – with an RMSD of just 1.4 Å between the two sites for all atoms in contact residues (on ligand and receptor). Very similar surface areas are also buried in the two sites (within 10%). Contrasting with this, one site in the asymmetric EREG-induced wild-type sEGFR dimer (left in Figure 1d) buries over 33% more surface area than the other (Freed et al., 2017) (2,891 Å^2^ compared with 1,938 Å^2^) – with a contact residue RMSD between the two sites of 3.0 Å (Figure S4d). The overall surface area buried upon EREG binding is also increased in the EREG-bound R84K sEGFR dimer compared with wild-type, suggesting an increase in affinity. Indeed, as shown in Figure 6b, the R84K mutation increases EREG-binding affinity of sEGFR by approximately 10-fold based on surface plasmon resonance (SPR) studies.

**FIGURE 6.**
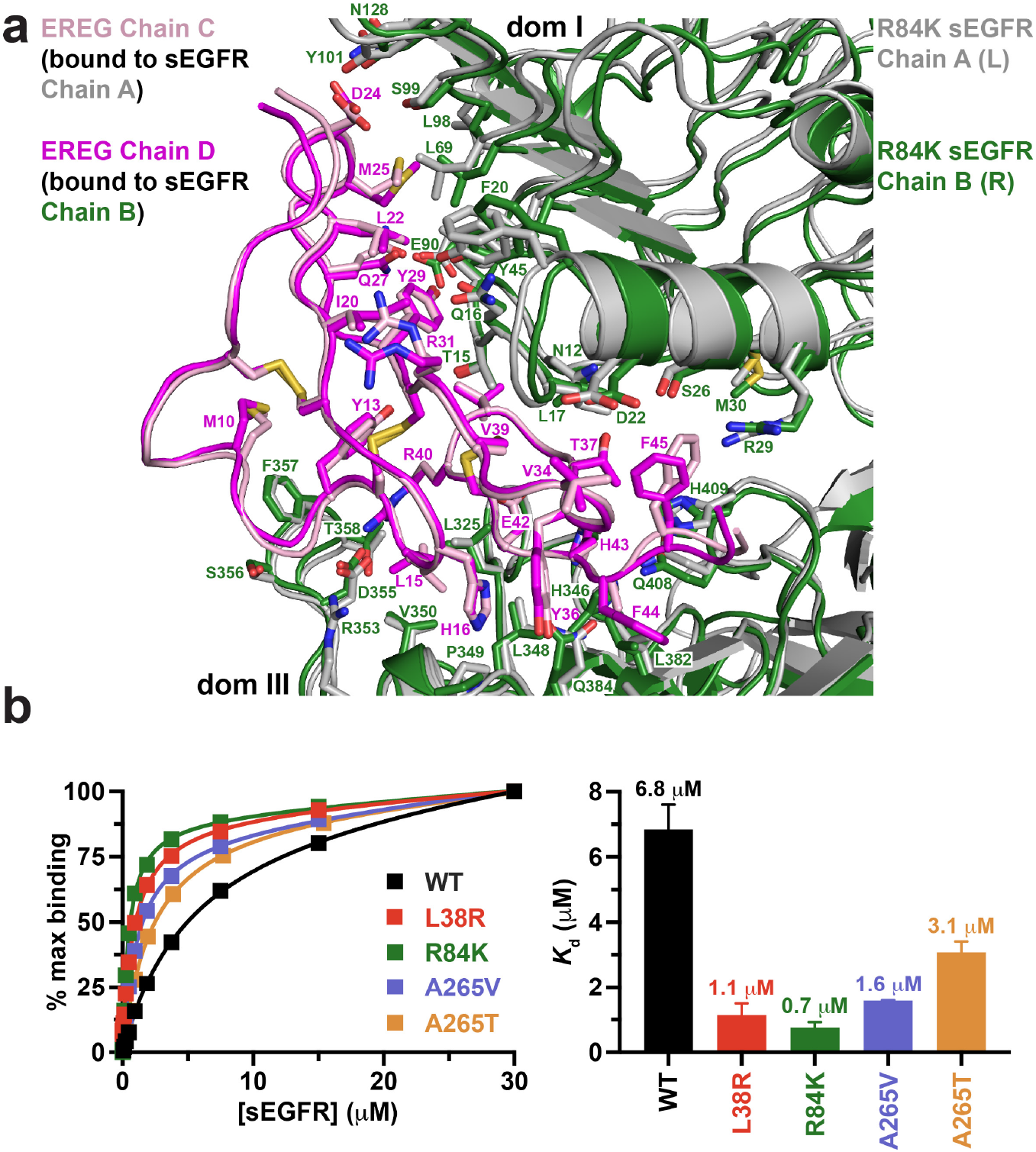
Similarity of the 2 ligand-binding sites in EREG-induced R84K sEGFR dimers. **a.** Comparison of the two EREG-binding sites in the symmetric dimer of R84K-mutated sEGFR dimer, overlaid by superposition of the ligand chains. Chain A of sEGFR (left in Figure 5a) is shown in grey ribbons, and chain B (right in Figure 5a) in dark green ribbons, and the respective bound ligands are colored pink and magenta. Side-chains involved in direct EREG/sEGFR contacts are shown and labelled. Those in the ligands superimpose very well (see Y13, H16, M25, Y29, for example), with a few exceptions (e.g. R31 and F45). Similarly, sEGFR side-chains in the binding sites overlay well, including D22, R29, Y45, E90 and S99 in domain I and D355, L348, F357, and H409 in domain III. Accordingly, the rmsd for all atoms in the 56 residues involved in ligand/receptor contacts (35 from sEGFR, 21 from EREG) is 1.4 Å. **b.** Comparison of EREG binding to different sEGFR variants as assessed using SPR (see Methods). This wild-type sEGFR construct bound to immobilized EREG in SPR studies with a *K*_d_ value of 6.84 ± 0.76 μM. GBM mutations in domain I increased ligand-binding affinity by 6.2 fold for L38R (p < 0.0001) and almost 10-fold for R84K (p < 0.0001), with *K*_d_ values respectively of 1.14 ± 0.36 μM and 0.75 ± 0.18 μM. Domain II GBM mutations increased ligand-binding affinity by 4.2-fold for A265V (p = 0.0008) and just ~2-fold for A265T (p < 0.03), with *K*_d_ values respectively of 1.58 ± 0.02 μM and 3.1 ± 0.3 μM. This smaller difference is consistent with the asymmetry retained in the A265V ligand binding sites. p values are for unpaired two-tailed Student’s t-tests. See also Figure S3.

The ability of the R84K mutation to equalize the two EREG-binding sites in a dimer and to increase EREG affinity suggests that it removes a barrier to ligand binding, and may diminish half-of-the-sites negative cooperativity seen in wild-type EGFR (Alvarado et al., 2010; Ferguson et al., 2020; Liu et al., 2012; Macdonald and Pike, 2008). As we described in detail for the *Drosophila* EGFR (Alvarado et al., 2010), and as also seems to apply to human EGFR (Liu et al., 2012), binding of a single ligand can promote formation of asymmetric sEGFR dimers in which autoinhibitory domain I/II interactions are broken in only one protomer (Figure 5e). Asymmetric dimerization is driven by contacts between N-terminal regions of domain II plus altered dimer arm docking (Alvarado et al., 2010; Freed et al., 2017) – together restraining domain II in the unliganded (right-hand) protomer in Figure 5e. When the second ligand binds to this dimer, it must ‘wedge’ apart the two ligand-binding domains (I and III) to drive formation of the symmetric dimer (upper part of Figure 5e). This requires disruption of autoinhibitory domain I/II interactions (formed by GBM-mutated residues) and the domain II dimer interface contacts – with resulting distortion of domain II (Figure S3d). Whereas this is readily achieved by high affinity ligands such as EGF and TGFα, low affinity EGFR ligands like EREG (Singh et al., 2016) cannot disrupt autoinhibitory domain I/II interactions – their binding energy apparently being insufficient to bend the restrained domain II to optimize dimer arm contacts. The consequent failure to wedge apart domains I and III in the right-hand protomer in Figure 5c (lower) requires that the second EREG binds instead to an unaltered asymmetric dimer using a compromized set of interactions (i.e. a remodeled binding site (Freed et al., 2017) – see below). The R84K mutation lowers this barrier to dimer ‘symmetrization’ by weakening the autoinhibitory domain I/II interactions so that the second ligand-binding event can more readily bennd domain II and symmetrize dimers. This is the origin of selective stabilization of dimers induced by low-affinity EGFR ligands by the R84K mutation. Weakening of the autoinhibitory domain I/II interactions also explains the enhanced ligand-binding affinity seen for R84K EGFR – both for EREG (Figure 6b) and for high-affinity ligands as we reported previously (Bessman et al., 2014a).

### Effects of other GBM mutations

The above analysis of the R84K mutation suggests that several other recurrent extracellular GBM mutations may influence EGFR activation in a similar way. We would predict that an L38R substitution breaks autoinhibitory domain I/II interactions in much the same way as R84K, directly disrupting the L38/F263 interaction seen in Figure 1e. Indeed, we find that EREG-induced sEGFR dimerization is selectively enhanced by the L38R mutation (Figure 3b, f) with an associated ~6-fold increase in EREG-binding affinity (Figure 6b). Interestingly, despite the participation of F263 in autoinhibitory domain I/II interactions, mutations at this site are not seen in GBM – consistent with the fact that F263 plays an integral role in forming the F263/Y275/R285 pocket that docks the dimer arm (Figure 5b-d). Accordingly, mutating F263 impairs function (Ogiso et al., 2002) rather than enhancing dimerization. Turning to other GBM mutations, R198 and R228 substitutions may disrupt domain I/II interactions with effects similar to those seen for L38R or R84K – with the caveat that R198 and R228 are most frequently substituted with cysteine, which might activate EGFR through aberrant disulphide crosslinking. We also investigated substitutions position 265, the most commonly mutated site. A265V or A265T mutations again selectively stabilize the EREG-induced sEGFR dimer (Figure 3d-f), but do so slightly less well than L38R or R84K. Both A265 substitutions also increase EREG binding affinity (Figure 6b), but again the effect is slightly smaller (by ~2-4 fold) than with L38R or R84K.

### A265V mutation reorients the dimer arm

The slightly reduced impact of A265 mutations on EREG-binding affinity and resulting sEGFR dimerization when compared with R84K prompted us to ask whether A265 substitutions stabilize sEGFR dimers through a distinct mechanism. Surprisingly, a 3.5 Å resolution crystal structure showed that the EREG-bound A265V sEGFR dimer remains asymmetric (Figure 7a, Table 1), contrasting with R84K. The autoinhibitory domain I/II interactions are retained in the right-hand protomer as in the EREG-induced wild-type sEGFR dimer (Figure S4a). The increased size of the side-chain at position 265 (going from A to V) leads to a slight (2-3 Å) displacement of the backbone C-terminal to this region (Figure S4b), since V265 still interacts with R84. The L38/F263 interaction is also retained, with F263 shifted in position by ~2 Å. The asymmetry evident in Figure 7a is clearly manifest in the difference between the two ligand-binding sites of the dimer, in a way that closely resembles the EREG-induced wild-type sEGFR dimer (Figures S4c and d). This suggests that negative cooperativity is retained, and may also explain why the A265V and A265T mutations have smaller influences on EREG-binding affinity than L38R or R84K (Figure 6b).

**FIGURE 7.**
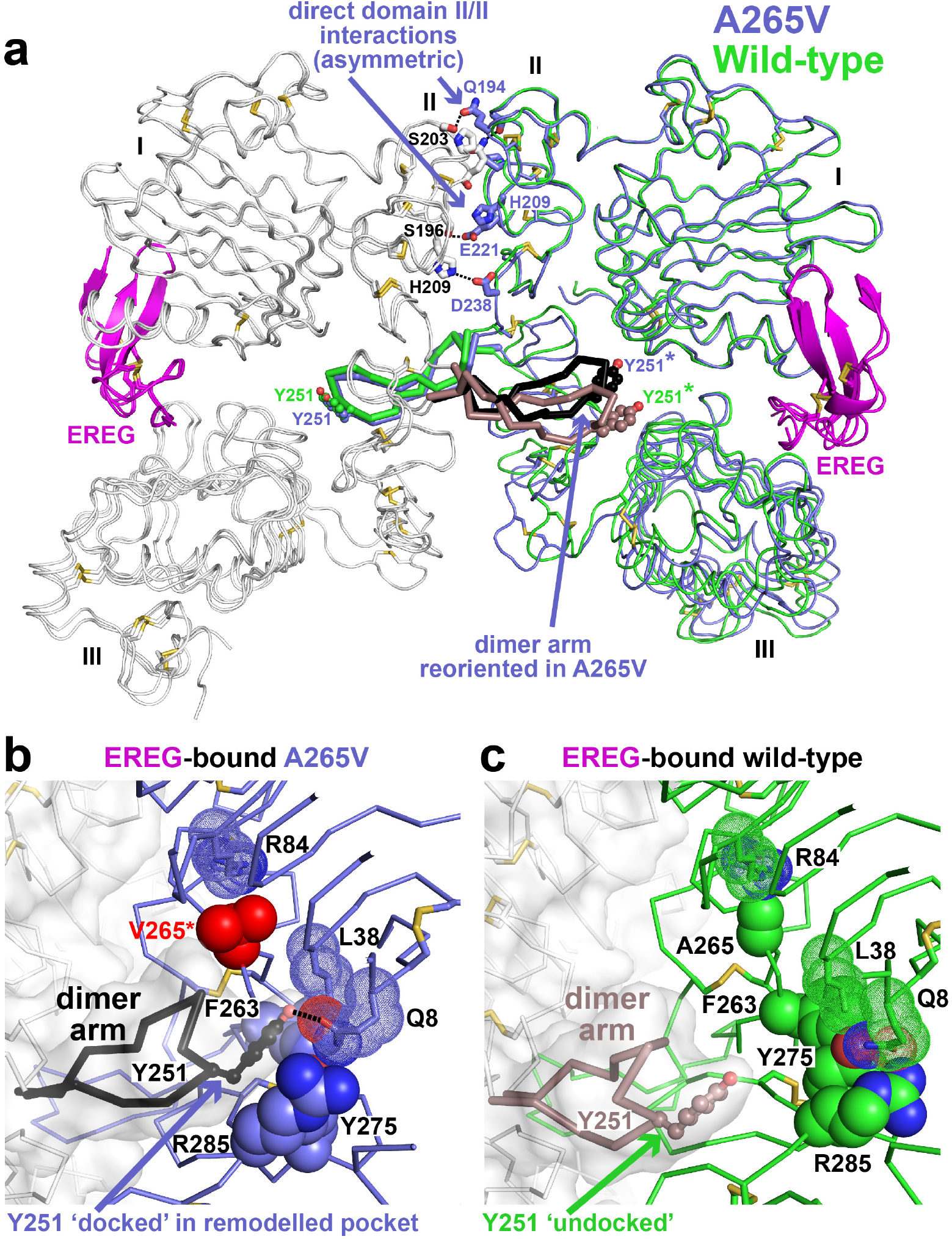
The A265V GBM mutation strengthens EREG-induced EGFR dimerization by optimizing dimer arm docking. **a.** Overlay of structures of EREG-induced dimers of wild-type sEGFR and A265V-mutated sEGFR. The left-hand molecule is grey in both dimers. The right-hand molecule is light green (wild-type) or slate blue (A265V). EREG is magenta. The key difference, as labelled, is reorientation of the dimer arm of the left-hand molecule in A265V sEGFR, where it is colored black, compared with salmon in the EREG-induced wild-type sEGFR dimer. The Y251 side-chain is labelled, as are the asymmetric domain II/domain II interactions seen in the asymmetric EREG-induced sEGFR dimer. **b.** Close-up showing the unique mode of docking of Y251 in the dimer arm in the A265V-mutated EREG-induced dimer. Y251 of the dimer arm (black) is docked into a binding site formed by the F263, Y275 and R285 side-chains (slate blue) with additional unique contributions from L38 and Q8 (from domain I) that do not contribute in the wild-type case (**c**). This minor remodeling of the docking site results when A265 is replaced with valine, leading to displacement of more C-terminal residues in domain II ‘downwards’ in the plane of the page (see Figure S4b). **c.** Close-up of undocked dimer arm (salmon) in EREG-induced wild-type sEGFR dimer as in Figure 5c. See also Figure S4.

With asymmetry and domain I/II autoinhibitory interactions retained, the A265V mutation must stabilize the EREG-induced dimer through a different mechanism than R84K. The overlay in Figure 7a draws attention to the substantial reorientation of the dimer arm in the A265V dimer (black) compared with its position the EREG-induced wild-type sEGFR dimer (salmon). Interestingly, this reorientation allows Y251 of the dimer arm from the left protomer to dock in a unique way onto a slightly remodeled pocket (Figure 7b) that includes, in addition to F263/Y275/R285, the side-chains of Q8 and L38 from domain I, which contribute directly to dimer arm docking in a way not seen in any other EGFR dimer (Q8 making a polar interaction with Y251 as shown in Figure 7b). This appears to be a direct consequence of the displacement of the backbone at position 265 mentioned above (Figure S4b), resulting from ‘growth’ of the side-chain at this position (still in contact with R84) from an alanine to a valine. The resulting movements of the F263, Y275 and R285 side-chains, plus direct contributions of Q8 and L38 effectively create the new binding site that accommodates the opposing dimer arm. The A265V variant thus simultaneously retains autoinhibitory domain I/II interactions and docks Y251 from the opposing molecule in a novel binding site not seen in wild-type sEGFR (Figure 7c).

Another characteristic of the EREG-induced dimer of A265V-mutated sEGFR is that it retains the asymmetric interface formed by the N-terminal part of domain II (Figure 7a). The interactions are driven by the same residues that stabilize the EREG-induced dimer of wild-type sEGFR (Freed et al., 2017) (Q194, S196, P204, H209, E221, and D238), which are also conserved in the asymmetric Drosophila EGFR dimer (Alvarado et al., 2010). This interface buries 634 Å^2^ of surface, compared with ~200 Å^2^ in the symmetric dimers (Freed et al., 2017). Alone, it can support only weak dimerization. When added to the remodeled dimer arm contacts shown in Figure 7b, however, this interface can support strong dimerization of the EREG/sEGFR complex.

## CONCLUSIONS

Our findings identify a primary ‘decision switch’ for biased agonism of EGFR, comprising autoinhibitory contacts between domains I and II of the receptor where GBM mutations are concentrated. A strongly-binding ligand like EGF or TGFα can overcome these autoinhibitory domain I/II contacts to optimize the strength of the resulting (symmetric) EGFR dimer. By contrast, weakly-binding ligands like EREG and epigen (EPGN) cannot overcome these interactions so readily. As a result, such ligands induce only weak, asymmetric dimers that signal with altered kinetics (Freed et al., 2017) – a feature of biased agonism that has now been observed in several receptor tyrosine kinases (Freed et al., 2017; Ho et al., 2017; Huang et al., 2017; Macdonald-Obermann and Pike, 2014; Zinkle and Mohammadi, 2018) and cytokine receptors (Kim et al., 2017; Moraga et al., 2015). GBM mutations that disrupt domain I/II autoinhibitory interactions in EGFR (R84K, A265V, A265T, L38R and likely others) remove this distinction, allowing EREG to induce strong EGFR dimers with characteristics similar to those normally driven by EGF or TGFα (or presumably betacellulin or HB-EGF (Singh et al., 2016)). Thus, GBM mutations effectively thwart the ability of EGFR to discriminate between its ligands.

The role and importance of EGFR in GBM present a substantial clinical puzzle (An et al., 2018a; Heimberger et al., 2005; Lee et al., 2006; Rutkowska et al., 2019; Vivanco et al., 2012). Despite a high frequency of EGFR amplification, deletion, or mutation in this cancer type, EGFR inhibitors have not been successful (Eskilsson et al., 2018), and the prognostic significance of EGFR aberration is complex (Eskilsson et al., 2018; Felsberg et al., 2017; Heimberger et al., 2005). The EGFR vIII variant is only weakly active (Rutkowska et al., 2019), and single amino acid substitutions seen in GBM do not promote strong ligand-independent dimerization (Bessman et al., 2014a; Orellana et al., 2019) – still requiring ligand for activation (Binder et al., 2018; Orellana et al., 2019). Rather than simply activating EGFR to promote tumor development as in lung cancer (Sharma et al., 2007), our data suggest that EGFR aberrations in GBM may instead alter the qualitative nature of EGFR signaling – as has also been suggested for EGFR vIII (Fan et al., 2013) – possibly by signaling to the microenvironment in a way that promotes tumor growth (An et al., 2018b). EREG is known to promote differentiation by activating EGFR in a plethora of cell types (Cao et al., 2020; Cui et al., 2019; Du et al., 2013; Freed et al., 2017; Jeong et al., 2020; Rizzi et al., 2013; Takahashi et al., 2003), and different EGFR ligands can promote preferential differentiation (Mukhopadhyay et al., 2013). One possibility suggested by our findings is that GBM mutations in EGFR ‘redirect’ responses of the receptor to activation by EREG so that they more closely resemble those seen with EGF. GBM mutations could thus impair the ability of EREG to promote normal differentiation of progenitor cells – effectively promoting dedifferentiation, which can induce gliomas (Friedmann-Morvinski et al., 2012) as seen with PLAGL2 amplification in GBM (Zheng et al., 2010).

What might our results mean for targeted therapy in GBM? Relatively small changes in conformation - arising from R-to-K or A-to-V mutations – can redirect EGFR signaling in a way that is relevant for cancer progression or maintenance. It should be possible to mimic small changes of this nature using conformation-specific or allosteric antibodies, which could function as biased agonists for therapeutic ‘correction’ of EGFR signaling – rather than applying frank inhibitors like kinase inhibitors or cetuximab. Importantly early studies demonstrated the possibility of generating EGFR antibodies that selectively impair binding of one ligand (TGFα) and not another (EGF) (Winkler et al., 1989), or that selectively bind only one or other ‘affinity class’ of EGFRs (Bellot et al., 1990; Defize et al., 1989).

The new element (and potential vulnerability) in the allosteric regulation of EGFR that we have identified here may also extend to other members of the EGFR family, and possibly to other receptors. Intriguingly, mutations in the corresponding domain I/II interaction region of ErbB3 are also found in cancer (Jaiswal et al., 2013), primarily in gastrointestinal tumors. More than 60% of the ErbB3 mutations reported in COSMIC (Tate et al., 2019) are in the ECR, with 28% (134 entries) at V104 (V85 in mature protein numbering: corresponding to R84 in EGFR) and 9% at G284 (G265 in mature ErbB3 numbering, which aligns structurally with A265 in EGFR). These mutations may selectively enhance heterodimerization of ErbB3 with EGFR or ErbB2 in response to certain ligands, which would further bias the complex network of ErbB receptor signaling and could rebalance proliferation versus differentiation. Mutations in the ErbB2 ECR (Greulich et al., 2012) found in tumors may also influence its heterodimerization preferences to influence signaling outcomes. The structural model for how these mutations influence signaling specificity suggested by our results with EGFR point to the potential value of a new therapeutic approach employing biologics (e.g. antibodies) as biased agonists, which would parallel efforts currently being pursued with small molecules for G-protein coupled receptors (GPCRs) (Lane et al., 2017; Smith et al., 2018) where biased agonism has long been known.

## EXPERIMENTAL PROCEDURES

### Protein Expression and Purification

DNA encoding the natural signal peptide and residues 1-501 of mature human EGFR, with a C-terminal hexahistidine tag, was subcloned into pFastbac1 (ThermoFisher Scientific) and recombinant baculovirus was generated using the Bac-to-Bac system. Q5 site-directed mutagenesis (New England BioLabs) was used to generate sEGFR variants harboring extracellular mutations. Protein expression was induced by baculovirus infection of 6-8 liter cultures of Sf9 cells in ESF921 medium (Expression Systems) at a density of ~2 × 10^6^ cells/ml. Conditioned medium was harvested 3-4 days post-infection, concentrated ~5-fold using a 10 kDa Sartocon Slice ECO Hydrosart cassette (Sartorius), and diafiltered against 4 volumes of 10 mM HEPES, pH 8.0, containing 150 mM NaCl (buffer A). The sample was then loaded onto 3 ml of Ni-NTA resin (Qiagen) by gravity at 4°C. After extensive washing with buffer A containing 10 mM imidazole, sEGFR proteins were eluted using an imidazole gradient ranging from 25 to 300 mM. Proteins were buffer exchanged into 25 mM MES, pH 6.0, containing 50 mM NaCl (buffer S) and loaded onto a Fractogel SO3^-^ cation exchange column (Millipore) that was subsequently developed using a gradient from 50 mM to 1 M NaCl in buffer S. sEGFR was eluted from the column during an isocratic step at 240 mM NaCl (24 mS/cm). Fractions containing sEGFR were pooled, concentrated, and purified further using a Superose 6 10/300 GL (Cytiva Life Sciences) equilibrated in buffer A. Protein purity was assessed using Coomassie-stained SDS-PAGE. Epiregulin was produced exactly as described (Freed et al., 2017), and human epidermal growth factor and TGFα were purchased (R&D Systems) and resuspended in buffer A.

### Small-Angle X-ray Scattering

SAXS data were recorded at 4°C on a Rigaku BioSAXS-2000nano 2D Kratky block camera system with a Rigaku 007HF rotating anode source and a Rigaku HyPix-3000 HPAD CCD detector, with 90 min exposures. Protein concentration was ~4 mg/ml (70 μM) in buffer A. Ligands were added at a 1.2-fold molar excess (84 μM), such that >90% saturation of sEGFR should be reached in each case. Data were reduced, and matched buffers were subtracted using BioXTAS RAW (Hopkins et al., 2017) to yield the corrected scattering profile – in which intensity (*I*) is plotted as a function of *q* (*q* = 4πsinθ/*λ*, where 2θ is the scattering angle). All samples were monodisperse as evidenced by linear Guinier regions (Figure 3). Values for *I*(0), the scattering intensity extrapolated to zero angle, were calculated from the Guinier region, where *q*\**R*_g_ < 1.4. Values were normalized by mass concentration of receptor protein, and were then divided by the *I*(0)/c value for an unliganded (monomeric) sEGFR sample collected on the same day to yield the fold-change in oligomeric state as described (Freed et al., 2017; Lemmon et al., 1997). The concentration-normalized intensity of forward scatter, *I*(0), is proportional to the weight-averaged molecular mass of molecules in a solution scattering sample (Lemmon et al., 1997). All SAXS experiments were repeated at least 3 times with different protein preparations.

### Covalent Crosslinking

Purified sEGFR proteins (5 μM) were incubated without ligand or with EGF (6 μM) or EREG (6 μM) in buffer A containing disuccinimidyl suberate (DSS) at 62.5 μM (ThermoFisher Scientific) for 30 min at room temperature, alongside untreated control. An aliquot (20 μl) was then mixed with 4X Pierce™ LDS sample buffer (ThermoFisher Scientific), boiled for 2 min, and analyzed by SDS-PAGE (4-12%) with Coomassie Blue staining. Gel images were quantitated using ImageJ (Figure S1).

### Crystallography

Crystals of sEGFR variants bound to epiregulin (EREG) were obtained using the hanging-drop method, mixing equal volumes of protein and reservoir solution and equilibrating this over reservoir solution at 21°C. For EREG:sEGFR^R84K^ crystals, a mixture of sEGFR^R84K^ (~8 mg/ml) and EREG (1.2-fold molar excess) was diluted 1:1 with reservoir solution containing 100 mM HEPES (pH 7.5), 12% PEG3350, with 3% (w/v) D-(+)-trehalose dihydrate. Crystals appeared within 3 days, and were cryoprotected in 100 mM HEPES (pH 7.5), 12% PEG3350, 7% glycerol, and 7% ethylene glycol. For EREG:sEGFR^A265V^ crystals, a mixture of sEGFR^A265V^ (~8 mg/ml) and EREG (1.2-fold molar excess) was diluted 1:1 with reservoir solution containing 1% w/v tryptone, 1 mM sodium azide, 50 mM HEPES (pH 7.0), 20% PEG3350. Crystals appeared within 3 days, and were cryoprotected in 100 mM HEPES (pH 7.5), 16% PEG3350, 7% glycerol, and 7% ethylene glycol.

Crystals of EREG-bound sEGFR^R84K^ diffracted to 2.9 Å resolution at the Advanced Photon Source (APS) GM/CA @ APS beamline, 23ID-B, and belonged to space group P2_1_2_1_2_1_ (Table 1). The asymmetric unit contained one 2:2 EREG:sEGFR^R84K^ dimer and 53% solvent. Similar crystals were also obtained using 10 mM spermine tetrahydrochloride instead of trehalose as additive that diffracted to 3.2 Å (PDB ID 7LFR) and gave the same conclusions (main chain atom root-meansquare deviation 0.5 Å between the two structures). Crystals of EREG-bound sEGFR^A265V^ diffracted to 3.5 Å resolution at GM/CA @ APS, and belonged to space group P2_1_ – with four 2:2 EREG:sEGFR^A265V^ dimers per asymmetric unit and 52% solvent. All data were collected at a wavelength of 1.033 Å. The two 2:2 EREG:sEGFR^A265V^ dimers overlay with a main chain atom root-mean-square deviation of 1.4 Å after refinement. Figures are generated with the B and C receptor chains (bound to ligand chains G and F respectively).

Datasets were integrated using XDS (Kabsch, 2010), and scaled using SCALA from the CCP4 program suite (CCP4, 1994). Structures were solved by molecular replacement using Phaser (McCoy et al., 2007), using the EGFR chains from an EREG-induced wildtype sEGFR dimer (PDB: 5WB7) (Freed et al., 2017) as search model. The resulting maps showed clear electron density for ligand in each binding site. Cycles of model building using Coot (Emsley and Cowtan, 2004) were alternated with rounds of refinement in Buster (Smart et al., 2012), Refmac (CCP4, 1994) or Phenix (Adams et al., 2010), employing composite omit maps generated using Phenix. TLS refinement (Winn et al., 2001) was employed in later stages, with anisotropic motion tensors refined for each of the receptor domains and ligand molecules. Final structures were refined using Phenix and validated with MolProbity (Chen et al., 2010) and wwPDB servers. Analysis of Ramachandran statistic for the final EREG:sEGFR^R84K^ model showed 93.3%, 6.3%, and 0.5% of residues in favored, allowed and disallowed regions respectively. For the EREG:sEGFR^A265V^, corresponding numbers were 93.8%, 6.0% and 0.2%.

### Sedimentation Equilibrium Analytical Ultracentrifugation (SE-AUC)

Ligand-induced dimerization of TGFα-bound sEGFR variants was analyzed by SE-AUC experiments using an XL-I analytical ultracentrifuge (Beckman) exactly as described (Dawson et al., 2005). Samples (at 2, 5, and 10 μM) of wild-type or mutated sEGFR in buffer A were analyzed both in the presence and in the absence of a 1.2-fold molar excess of TGFα, used (rather than EGF) because it contributes very little to absorbance at 280 nm, having just one tyrosine and no tryptophans. Radial *A*_280_ data were collected at 20°C at 6,000, 9,000, and 12,000 rpm using an AnTi 60 rotor. The resulting nine datasets (three concentrations at three speeds) were fit to a model describing simple dimerization of a 1:1 sEGFR/TGFα complex, assuming that all sEGFR was saturated with TGFα and that TGFα does not contribute significantly to *A*_280_: *A_r_* = *A*_0_exp[*H · M*(*r*^2^ - *r*_0_^2^)] + *A*_0_^2^ · *K_a_exp*[*H · 2M*(*r*^2^ - *r*_0_^2^)], where *A_r_* is the absorbance at radius *r, A*_0_ is the absorbance at the reference radius *r*_0_, *M* is the molecular weight of the 1:1 sEGFR/TGFα complex (the sum of the measured monomeric sEGFR and TGFα molecular weights), *H* is the constant [(1 – ∇ ρ)*ω*^2^]/*2RT*, ∇ is the partial specific volume (estimated at 0.71 ml/g), ρ is the solvent density (1.003 g/ml), *ω* is the angular velocity of the rotor (radians/sec), *R* is the gas constant, *T* is the absolute temperature, and *K*_a_ is the fitted parameter corresponding to the equilibrium constant for dimerization of the 1:1 sEGFR/TGFα complex. The fitted *K*_a_ value is converted to the dissociation constant *K_d_* (*K_d_* = *1/K_a_*) reported in Figure S2a by using the calculated extinction coefficient for the 1:1 sEGFR/TGFα complex. At least three independent groups of experiments were performed (except L38R) and fit for each mutated protein. Estimated *K_d_* values are quoted the mean ± standard deviation of estimates from individual experiments. Data fitting used HeteroAnalysis (Version 1.1.0.58), from the U. Conn Biophysics Facility.

### Surface Plasmon Resonance (SPR)

SPR analysis of ligand binding was performed using a Biacore 3000 instrument exactly as described (Ferguson et al., 2000). EREG was immobilized on a CM5 sensorchip (Cytiva Life Sciences) using amine coupling, to a final level of ~2,500 Response Units. Purified sEGFR variants were injected onto the sensorchip at a variety of concentrations at 5 μl/min for 8 min (sufficient for binding to reach steady state) in degassed 10 mM HEPES (pH 7.5), 150 mM NaCl, 3 mM EDTA and 0.005% Surfactant P-20 at room temperature. Between injections, the sensorchip surface was regenerated using a 20 μl injection of 10 mM sodium acetate (pH 5.0) containing 1 M NaCl. The final steady-state signal was background-corrected by subtracting the signal obtained with a control surface. To estimate receptor/ligand affinities, steady state SPR signal values were plotted against [sEGFR] and fitted to a simple single-site saturation-binding model.

### Cell signaling and proliferation studies

IL-3 dependent murine Ba/F3 cells were cultured in RPMI 1640 (LifeTech 11875-093) supplemented with 10% FBS, 1 mM pyruvate, 10 mM HEPES, 1 ng/ml IL-3 (PeproTech #213-13) and PenStrep. Fully-haploid engineered HAP1 (eHAP) human cells (Essletzbichler et al., 2014) were obtained from Horizon (now PerkinElmer), and were cultured in complete IMDM medium (ThermoFisher # 12440-053) containing 10% FBS and 100 U/ml penicillin, with 100 μg/ml streptomycin. Transfections of full-length human EGFR (wild-type or noted variants) into Ba/F3 or eHAP cells were performed by electroporation using a Nucleofector 2b device (Lonza) as described previously (Kiyatkin et al., 2020). Transfected cells were selected for 2 weeks in G418-containing medium. Expression levels of EGFR variants in eHAP cells were confirmed by Western using anti-EGFR (R&D AF231). Stably transfected Ba/F3 cells were sorted by flow cytometry to select cells with similar expression levels of wild-type or mutated EGFR.

For Ba/F3 cell proliferation studies, cells were plated in triplicate in 96-well flat bottom plates in starvation medium without FBS or IL-3. Cells were either left untreated or were stimulated with EGF (100 ng/ml), EREG (50 μg/ml) or IL-3 (2 ng/ml) for 72 hours. Viable cell numbers were detected with the CyQuant Direct assay (Invitrogen #C35011) measuring fluorescence signals (excitation 485 nm, emission 528 nm) with a BioTek Synergy plate reader.

For cell signaling studies, eHAP cells were serum-starved overnight and either left unstimulated or stimulated with 100 ng/ml EGF for the indicated time intervals. Total cell lysates were prepared and analyzed by Western blotting as described previously (Kiyatkin et al., 2020). Chemiluminescence signals for phosphorylated EGFR (pY845: CST #2231) and ERK1/2 (pT202/pY204: CST #9106), were visualized and quantitated using a Kodak Image Station 440CF (Kodak Scientific). Blotting for Grb2 (Santa Cruz sc-255) was used as a loading control as described (Aksamitiene et al., 2015).

### Reporting summary

Further information on research design is available in the Nature Research Reporting Summary linked to this paper.

### Data availability

Atomic coordinates and structure factors have been deposited in the Protein Data Bank (PDB) under accession codes 7LEN and 7LFS. Source data are provided with this paper.

## ACKNOWLEDGEMENTS

We thank members of the Lemmon and Ferguson laboratories for discussions and comments on the manuscript. This work was supported by NCI grant R01-CA198164 (to M.A.L. and K.M.F.). Crystallographic data were collected at GM/CA@APS, funded by NCI (ACB-12002) and NIGMS (AGM-12006). The Eiger 16M detector at GM/CA-XSD was funded by NIH grant S10 OD012289. This research also used resources of the Advanced Photon Source, a U.S. Department of Energy (DOE) Office of Science User Facility operated for the DOE Office of Science by Argonne National Laboratory under Contract No. DE-AC02-06CH11357.

## AUTHOR CONTRIBUTIONS

C.H., K.M.F., and M.A.L. conceived the project. C.H. performed all protein production, purification, crosslinking, analytical ultracentrifugation and binding studies, with assistance from C.A.L. C.H. and S.E.S. performed all SAXS studies. C.H., K.M.F., C.A.L. and S.E.S. performed and/or interpreted all crystallographic analysis. A.K. performed cell signaling studies. M.A.L. and K.M.F. supervised the project. C.H. and M.A.L. drafted the manuscript, and all authors commented on the manuscript.

