## Supplementary Figures 1-4 for "Glioblastoma mutations impair ligand discrimination by EGFR"

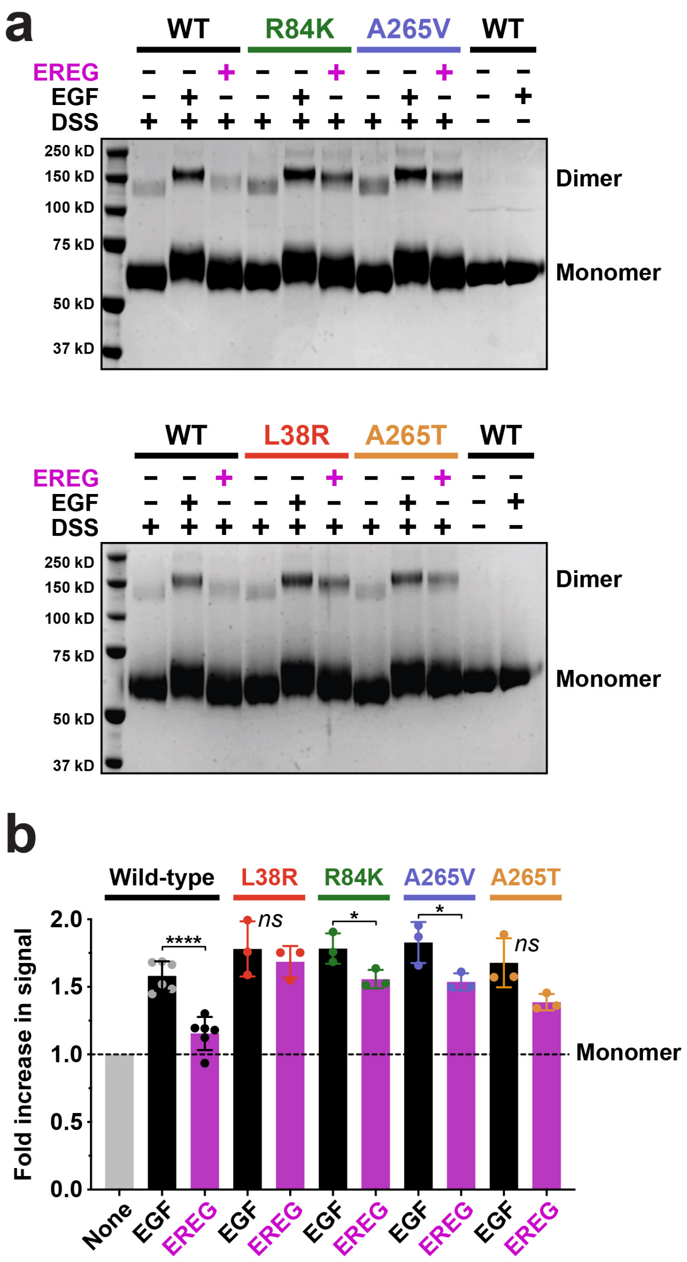


**FIGURE S1 – Related to Figure 3.**

**Chemical cross-linking studies of EREG-induced sEGFR dimerization**

**a.** Different sEGFR variants at 5 μM were incubated alone or with the noted ligand (EGF or EREG) at 6 μM, and subjected to 62.5 μM DSS for 30 min as described in Experimental Procedures. Samples were then subjected to SDS-PAGE and Coomassie Blue staining. Dimer and monomer bands are marked, with representative dimer bands shown in Figure 3 (note shift in monomer band upon cross-linking to ligand).

**b.** Quantitation of data in **a.** with additional repeats (at least 3) for each variant. For wild-type sEGFR, EGF induces substantially more dimerization than EREG ($p < 0.0001$), whereas the difference is not significant for L38R or A265T, and only just reaches statistical significance for R84K ($p = 0.04$) and A265V ($p = 0.04$). p values are for unpaired two-tailed Student's t-tests.

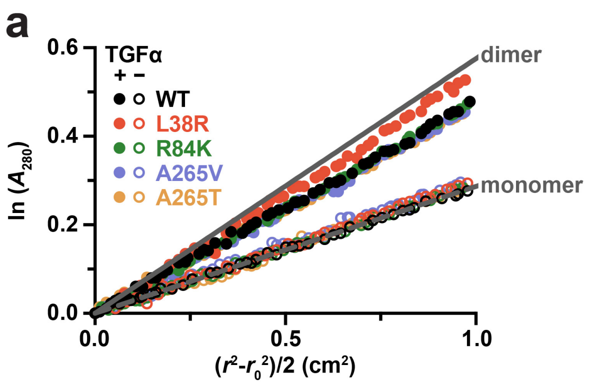


| Mutation | Dimerization $K_d$ |
|---|---|
| None | 1.26 +/- 0.30 μM |
| L38R | 0.65 μM |
| R84K | 1.58 +/- 0.32 μM |
| A265V | 3.45 +/- 0.63 μM |
| A265T | 3.60 +/- 0.66 μM |

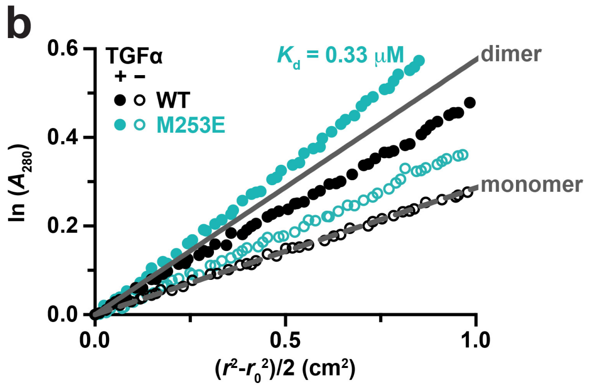


**FIGURE S2 – Related to Figure 5.**
**SE-AUC studies of TGFα-induced sEGFR dimerization**

**a.** The noted sEGFR variants were subjected to SE-AUC as described in Methods, with or without a 1.2-fold excess of TGFα. Representative data are shown for 10 μM sEGFR at 6,000 rpm, with the natural logarithm of the absorbance, ln($A_{280}$), at radial distance $r$ plotted against ($r^2 - r_0^2$)/2. This transformation of the data gives a straight line for a single species, with slope proportional to molecular weight. Expected data for pure monomer and pure sEGFR:TGFα dimer are shown as dotted and solid grey lines (marked). Data points are color coded for the different variants as described in the legend, with filled circles representing data with added ligand and open circles without. No dimerization was seen in the absence of ligand for any variant under these conditions, consistent with the SAXS studies shown in Figure 2 and our previous work (Bessman et al., 2014). $K_d$ values for each sEGFR:TGFα complex are listed below the graph, determined by global fit of SE-AUC data as described previously (Dawson et al., 2005) and in the Methods. Mean values ± S.D. for at least 3 biological repeats are reported for all cases except L38R (where n = 1). Whereas the R84K variant dimerizes with essentially the same $K_d$ as wild-type following TGFα binding, the A265V and A265T variants actually dimerize slightly more weakly (p = 0.006 and 0.005 respectively, for unpaired two-tailed Student's t-tests).

**b.** SE-AUC analysis of sEGFR harboring the M253E mutation seen in lung cancer (Tate et al., 2019; Yu et al., 2017) (but not seen in GBM), and comparison with wild-type. Unlike the GBM variants, M253E-mutated sEGFR dimerizes constitutively, being substantially dimeric in the absence of ligand. TGFα-bound M253E sEGFR also appears to form species larger than dimers, with an estimated $K_d$ in the range of 0.33 μM. M253E-mutated sEGFR was used at 10 μM, and the sample was spun at 6,000 rpm.

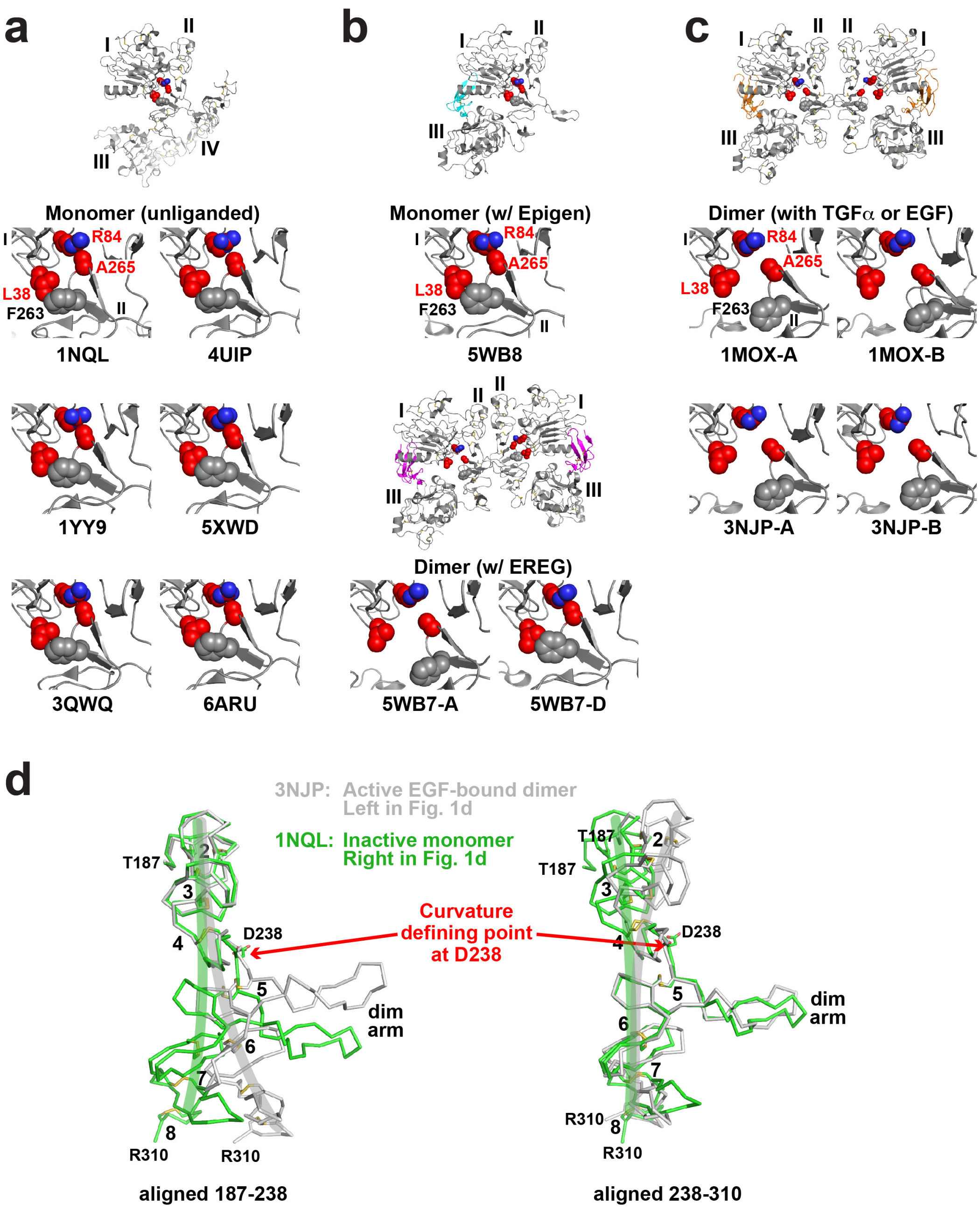


**FIGURE S3 – Related to Figure 6.**
**Autoinhibitory domain I/II interactions in different sEGFR structures**

**a.** Disposition of key GBM-mutated residues (L38, R84 and A265, colored red) in 'inactive' configurations of the EGFR extracellular region. The positions of these side-chains are shown in monomeric tethered forms of sEGFR (Ferguson et al., 2003; Lee et al., 2015; Li et al., 2005; Matsuda et al., 2018; Ramamurthy et al., 2012). In each case, the side-chain of R84 directly contacts that of A265, and the L38 side-chain is in van der Waal's contact with that of F263 (grey spheres: not mutated in GBM). These represent autoinhibitory interactions between domains I and II as described in the text and pointed out in our previous studies (Alvarado et al., 2009). This configuration is represented as a filled red star in the cartoons in Figure 3.

**b.** Importantly, the autoinhibitory R84/A265 and L38/F263 interactions are retained in the ligand-bound monomer seen when epigen binds to wild-type sEGFR (PDBID: 5WB8) (Freed et al., 2017). In the asymmetric EREG-induced dimer of wild-type sEGFR (5WB7: Figure 1d), the autoinhibitory interactions are retained in the right-hand molecule, but are lost in the left-hand molecule as described in the text.
**c.** As expected for autoinhibitory interactions, the R84/A265 and L38/F263 interactions are broken in the 'active' symmetric dimers of sEGFR induced upon activation with TGFα (PDBID: 1MOX) (Garrett et al., 2002) or EGF (PDBID: 3NJP) (Lu et al., 2010; Ogiso et al., 2002). This configuration is represented as open red stars in the cartoons in Figure 3.
**d.** Comparison of the bend in domain II in inactive and monomeric forms of sEGFR (light green) and active (dimeric) forms (grey) – colors corresponding to those used for sEGFR chains in Figure 1d. The structures of unliganded monomeric sEGFR (PDBID: 1NQL) (Ferguson et al., 2003) and an EGF-induced wild-type sEGFR dimer (PDBID: 3NJP) (Lu et al., 2010) were used. Only residues 187-310 of domain II are shown. In the left-hand panel, the two structures are overlaid using residues 187-238 as reference. In the right-hand panel, residues 238-310 are used as reference. This analysis reveals that the two structure differ by a bend at residue D238 (marked as 'Curvature defining point'), as mentioned in the main text. The approximate direction of curvature is shown by green and grey brush strokes on each structure. The dimer arm is labelled, as are disulphide-bonded modules 2-8 of domain II (Ferguson et al., 2003). This figure is based on one by Ferguson (Ferguson, 2008).

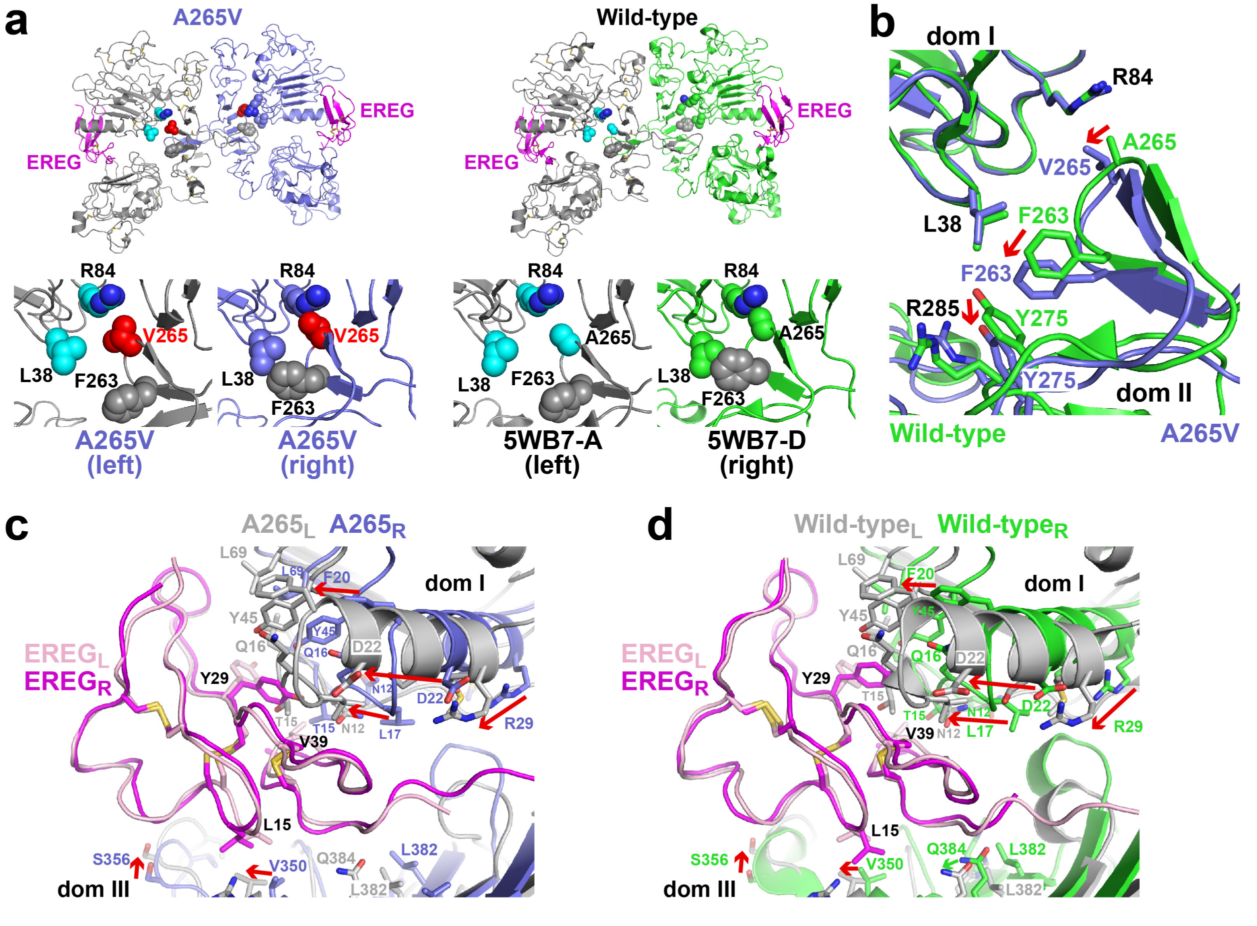


**FIGURE S4 – Related to Figure 7.**

**Structural features of the asymmetric dimer of A265V-sEGFR induced by EREG**

**a.** The asymmetric A265V-mutated (left) and wild-type (right) sEGFR dimers induced by EREG are compared, with disposition of the autoinhibitory domain I/II residues shown in the lower panels. Colors parallel those used in Figure 3a, with the right-hand molecule colored slate blue for A265V and light green for wild-type sEGFR. Side-chain contacts between residues at positions 38 and 263 and between residues 84 and 265 are retained in the right molecule but not the left in each case. V265, corresponding to the GBM mutation, is colored red.

**b.** Structural consequences of the A265V mutation in the domain I/II interface of the EREG-induced sEGFR dimer. The right-hand side of the EREG-bound wild-type and A265V-mutated sEGFR structures shown in **a** are superimposed using domain I as reference, with wild-type sEGFR shown in light green and A265V sEGFR in slate blue. Replacement of A265 with a valine causes a displacement of the Cα position for residue 265 by ~1.2 Å (red arrow), which is propagated to a shift in position of F263 by ~2.5 Å (red arrow). As a result of the small displacement in domain II constituents beyond this position, the locations of Y275 and R285 – which provide the docking side for the dimer arm Y251 residue in Figure 4b – are altered, allowing remodeling of this binding site to enhance dimerization strength as described in the text.

**c.-d.** Comparison of the binding sites on the two sides of the EREG-induced dimer for A265V-mutated sEGFR (**c**) and wild-type sEGFR (**d**), illustrating that the differences seen between the two sites in the asymmetric wild-type dimer (Freed et al., 2017) are retained in the A265V variant despite stronger dimerization and slightly stronger ligand binding. The regions corresponding to the two ligand-binding sites are superimposed with EREG as the reference. The left-hand molecule is colored grey in each case, and the right-hand molecule slate blue in **c** (A265V) and light green in **d** (wild-type). The pink ligand is bound to the left-hand (grey) sEGFR molecule, and the magenta ligand is bound to the right-hand sEGFR molecule. For clarity, ligand side-chains that are not substantially different in orientation are omitted – the exceptions being L15, Y29, and V39, which are consistently reoriented between the two sites. Contact side-chains in the receptor are shown, illustrating their substantial displacement with respect to the ligand in the two sites. Examples include the ~10 Å displacement of D22 and R29, and ~7 Å displacements of L17 and F20 that are marked by red arrows in domain I. These changes are essentially the same in A265V and wild-type. Shifts in domain III are generally smaller, but are essentially the same in A265V and wild-type sEGFR. Thus, the compromised binding to the right-hand molecule previously reported (Freed et al., 2017) is fully retained in the A265V variant despite stronger dimerization.
